# *MYBPHL* nonsense mutations have poor sarcomere binding, are degraded, and cause abnormal contraction

**DOI:** 10.1101/2024.07.01.601577

**Authors:** Alejandro Alvarez-Arce, Geena E Fritzmann, Hope V Burnham, Kelly N Araujo, Alexandra Pena, Lucas Wittenkeller, David Y. Barefield

## Abstract

Heart function depends on the cardiomyocyte contractile apparatus and proper sarcomere protein expression. Mutations in sarcomere genes cause inherited forms of cardiomyopathy and arrhythmias, including atrial fibrillation (AF). Recently, a novel sarcomere component, myosin binding protein-H like (MyBP-HL) was identified. MyBP-HL is mainly expressed in cardiac atria and shares homology to the last three C-terminal domains of cardiac myosin binding protein-C (cMyBP-C). The *MYBPHL* R255X mutation has been linked to atrial enlargement, dilated cardiomyopathy, and atrial and ventricular arrhythmias. Similar nonsense mutations in *MYBPC3* result in no myofilament incorporation and a rapid degradation of the truncated protein and are highly associated with development of hypertrophic cardiomyopathy. However, the *MYBPHL* R255X mutation occurs too frequently in the human population to be highly pathogenic. We sought to determine whether all *MYBPHL* nonsense mutations lead to impaired MyBP-HL sarcomere integration and degradation of the mutant protein, or if the *MYBPHL* R255X mutation has a different consequence. We mimicked human *MYBPHL* nonsense mutations in the mouse *Mybphl* cDNA sequence and tested their sarcomere incorporation in neonatal rat cardiomyocytes. We demonstrated that wild type MyBP-HL overexpression showed the expected C-zone sarcomere incorporation, like cMyBP-C. Nonsense mutations showed defective sarcomere incorporation. We demonstrated that wild type MyBP-HL and MyBP-HL nonsense mutations were degraded by both proteasome and calpain mechanisms. Additionally, we observed changes in contraction kinetics and calcium transients in cells transfected with MyBP-HL nonsense mutations compared to MyBP-HL full length. Together, these data support the hypothesis that *MYBPHL* nonsense mutations are largely similar.

**Short summary:** Premature stop mutations in myosin binding protein H-like prevent sarcomere incorporation of the translated protein. Overexpression of truncating mutants causes contractile defects in neonatal rat cardiomyocytes. These effects occur regardless of the location of the premature stop along the protein.

## Introduction

Heart function relies significantly on the proper regulation of the cardiomyocyte contractile apparatus by sarcomere protein expression. Genetic mutations within the genes encoding these proteins can result in hereditary cardiomyopathies and arrhythmias (Ludhwani and Wieters, 2022). The most common cardiomyopathies are hypertrophic (HCM) occurring in 1:500 people (Maron et al., 2022a) and dilated (DCM) that occurs in 1:250 people (Marschall et al., 2019; Hershberger et al., 2021). About 60% of HCM cases are caused by sarcomere gene mutations, while about 30% of DCM is related to sarcomere gene mutations (Maron et al., 2022b; Barefield et al., 2023a). Atrial fibrillation is highly associated with both HCM and DCM and represents the most common arrythmia (Dragasis et al., 2022).

A myosin binding protein H-like (MyBP-HL) nonsense variant, R255X is associated with ventricular conduction system abnormalities, atria enlargement, arrhythmia, and DCM in humans. A mouse with an *Mybphl* null allele produced atrial and ventricular dilation, systolic dysfunction, premature ventricular contractions, and cardiac conduction system dysfunction. Importantly, the R255X variant promotes an improper myofilament incorporation (Barefield et al., 2017). Myosin binding protein H-like (MyBP-HL) is a ∼40 kDa myofilament protein that is highly expressed in the atria with a small subset of expression near the ventricular conduction system (Barefield et al., 2022). MyBP-HL is composed of three domains: HL1 immunoglobulin (Ig) domain, HL2 fibronectin-III (FN3) domain, and HL3 Ig domain which regulate MyBP-HL sarcomere incorporation.

Cardiac myosin binding protein C (cMyBP-C) is a ∼140 kDa sarcomere accessory protein encoded by the *MYBPC3* gene. cMyBP-C is a multi-modular structural protein component of the sarcomere, comprised of Ig domains (C1-C5, C8, C10) and FN3 domains (C6, C7, C9). MyBP-HL domains HL1-HL2-HL3 share similarities in sequence and structure to last three C-terminal domains from cMyBP-C (C8-C9-C10) (Barefield et al., 2017). These three C-terminal domains are required for proper sarcomere incorporation in cMyBP-C (Welikson and Fischman, 2002). Nonsense mutations in *MYBPC3* are the most common cause in HCM (Fernandez Suarez et al., 2023). Importantly, *MYBPC3* nonsense mutations cause improper myofilament incorporation and degradation of the truncated protein (van Dijk et al., 2009). Nonsense mutations lead to a reduction in cMyBP-C content in the myocardium and improper actomyosin-crossbridge regulation, causing an increase in myofilament sliding velocities and calcium sensitivity $. We have recently shown that reduction in MyBP-HL levels causes a concomitant increase in cMyBP-C levels in atrial sarcomeres, suggesting that nonsense mutations in *MYBPHL* may cause increased levels of atrial cMyBP-C and lead to dysfunction. Notably, the population frequency of the R255X variant is 9.21^e-4^ (Chen et al., 2024)$, too high to be a fully penetrant disease-causing gene. While the role of atrial contractile dysfunction in the development of overall cardiac dysfunction remains unclear, we wanted to investigate whether the R255X variant behaves differently than the other observed *MYBPHL* nonsense mutations.

In this work, we overexpressed mouse MyBP-HL in neonatal rat myocytes (NRM’s) to determine MyBP-HL nonsense mutation sarcomere incorporation. We used both atrial and ventricular NRMs (NRAM and NRVM, respectively) to model a system with no endogenous MyBP-HL (NRVMs) and one with endogenous rat MyBP-HL in the atria. We identify that overexpression of truncated MyBP-HL does cause alterations in sarcomere structure and dysregulation of cardiomyocyte function. We conclude that all *MYBPHL* nonsense mutations have a similar effect, causing MyBP-HL to not incorporate into the sarcomere.

## Methods

### NRAM/NRVM isolation

All animals were housed and sacrificed in accordance with Loyola University Chicago’s Institutional Animal Care and Use Committee (IACUC) in adherence to the US National Institutes of Health Guide for Care and Use of Laboratory Animals. Neonatal rat hearts were isolated from 0 – 1-day old Sprague-Dawley pups (Charles River Laboratories). Briefly, hearts were rinsed twice in ice-cold, filtered Krebs-Henseleit Buffer. Both atrial and ventricular myocytes were isolated separately as described previously (Martin et al., 2021).

### Plasmid design

We used the information reported by gnomAD v2.1.1 about human *MYBPHL* nonsense mutations, then mimicked *MYBPHL* nonsense mutations using the mouse *MYBPHL* (NM_026831.1) sequence. We designed specific primers (Table 1) for site-directed mutagenesis to generate nonsense mutations in mouse *Mypbhl* using Q5 Hot Start High-Fidelity (New England Biolabs, Cat No: M0494S). Plasmids were sequenced by Sanger reaction (Azenta Life Sciences, Burlington, MA). Transfection quality plasmid was prepared for all plasmids with confirmed nonsense mutations using an endo-free maxi prep kit (Qiagen, Cat. No: 12362).

**Table 1.**
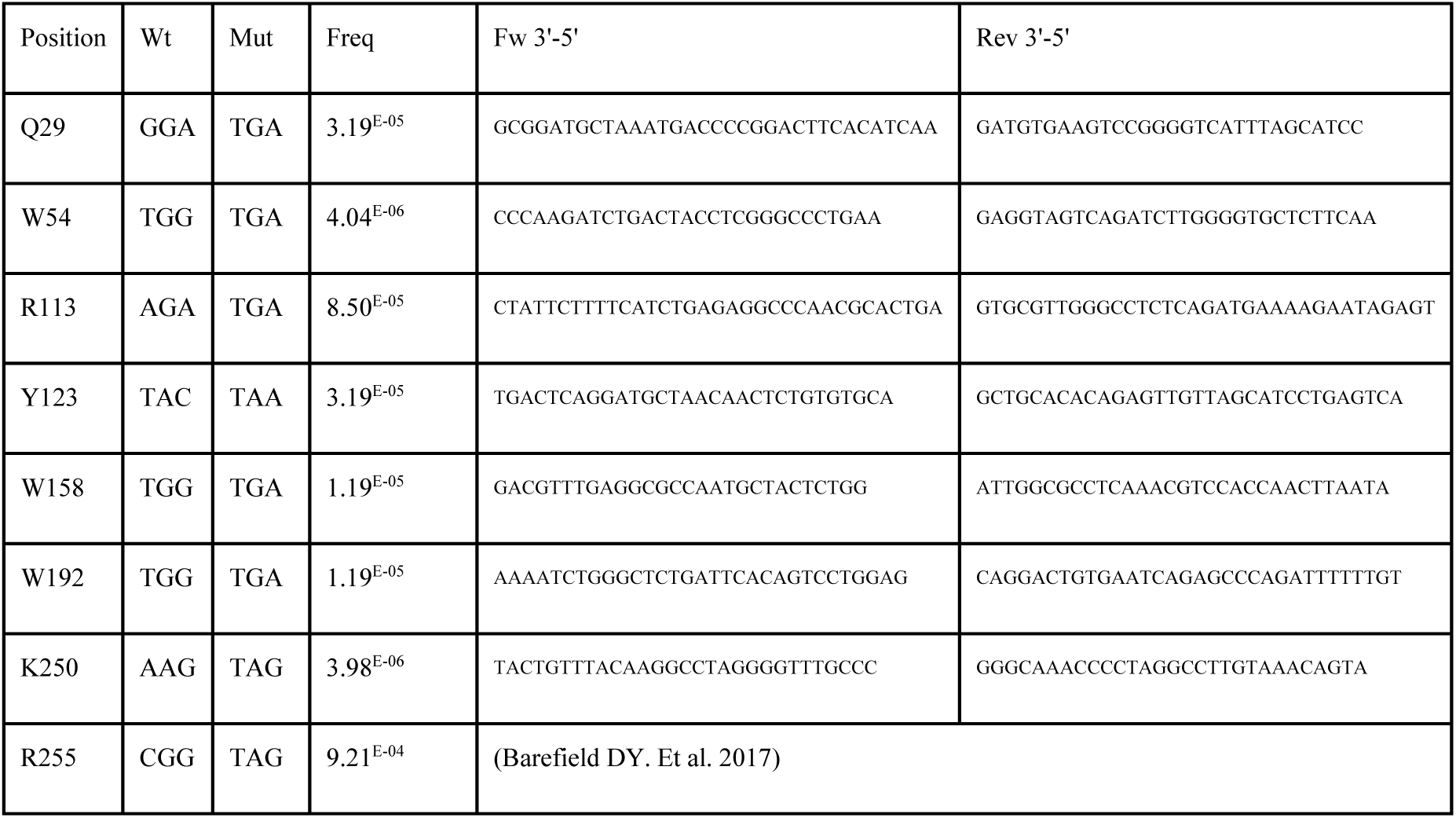
MyBP-HL nonsense mutations primers. We designed primers to modify the mouse *Mybphl* full length sequence through direct mutagenesis and mimic the MyBP-HL nonsense mutations reported in gnomAD.

### Immunoblotting

To evaluate protein expression of MyBP-HL nonsense mutations in NRM’s and HEK, cells were seeded on 6-well plates coated with 0.1% gelatin (Sigma Aldrich, Cat. No: G2500-100G) at 1×10^6^ cells per well. Cells were then transfected with 2.5 mg of full length MyBP-HL or MyBP-HL nonsense mutations using Lipofectamine 3000 (Thermofisher, Cat. No: L3000008). Calpastatin (1 µM, Sigma, Cat. No: 208902) and/or epoxomicin (100 nM, AbCam, Cat. No: ab144598) were added to the culture 24 hours before lysing. NRM’s were rinsed with 1x PBS and then lysed in RIPA buffer (50 mM Tris HCl, 150 mM NaCl, 1.0% (v/v) NP-40, 0.5% (w/v) sodium deoxycholate, 1.0 mM EDTA, 0.1% (w/v) SDS, and 0.01% (w/v) sodium azide, pH 7.4) containing protease inhibitor cocktail (Millipore Sigma, Cat. No: 11836170001). Protein concentration was determined using a Pierce BCA Protein Assay Kit (ThermoFisher, Cat. No: 23227) and standardized. Laemmli buffer (0.75 mM Tris-HCl; pH 8.8, 5% SDS, 20% glycerol, 0.01% bromophenol blue) supplemented with 5% β-mercaptoethanol, was added to the lysates and samples were heated at 75 °C for 5 minutes. Samples were resolved by 10% SDS-PAGE, and transferred onto PVDF membranes (BioRad, Cat. No: 1620177). Membranes were blocked with tris buffered saline (TBS) with 1% casein blocker (Biorad, Cat. No: 1610782) for one hour. Primary antibody against mouse MyBP-HL (1:2500) was incubated in blocking buffer overnight at room temperature (Barefield et al., 2022). Secondary goat anti-mouse (Jackson ImmunoResearch, Cat. No: 115-035-003) was probed in blocking buffer for one hour. Membranes were incubated with HRP substrate Clarity Western ECL (Biorad, Cat no: 1705062) for 10 sec, and luminescence was captured with an Azure Biosystems c600 imager. Densitometry analysis was performed using Fiji software, and data were analyzed with Prism8 software (GraphPad, San Diego, CA).

### Immunofluorescence staining and confocal microscopy

To confirm if MyBP-HL mutations were incorporated into the sarcomere of NRM’s, 1.75 x 10^5^ myocytes were plated on 12 mm glass coverslips (Fisher Scientific, Cat. No: 12-541-000) coated with GelTrex Membrane matrix (ThermoFisher, Cat. No: A1413202). Cells were transfected with 2.5 µg of full length MyBP-HL or MyBP-HL nonsense mutations using Lipofectamine 3000 (ThermoFisher, Cat. No: L3000008). Cells were fixed with 4% PFA (ThermoFisher, Cat. No: 50-980-487) and permeabilized with 0.25% Triton X-100 (Millipore Sigma, Cat. No:X100). Cells were blocked with 20% FBS, 0.1% Triton in PBS. Primary antibodies against MyBP-HL (Barefield, 2022; 1:2500, (Barefield et al., 2022)) and cMyBP-C (Santa Cruz Tech, E7; Cat. No: sc-137180; 1:2500) were incubated in blocking buffer overnight at 4 °C. Secondary antibodies donkey anti-Mouse IgG Alexa Fluor 488 (ThermoFisher Cat. No: A21202; 1:2500) and donkey anti-rabbit IgG Alexa Fluor 568 (ThermoFisher Cat. No: A10037; 1:2500) were incubated in blocking buffer for one hour at room temperature. Nucleus staining was done with 1x Hoechst for 10 minutes. Coverslips were mounted using ProLong Gold (ThermoFisher, Cat. No: P36930). Images were obtained using a Zeiss 3i microscope, connected to a W1 Confocal Spinning Disk system (Yokogawa) equipped with Mesa field flattening (Intelligent Imaging Innovations), a motorized X,Y stage (ASI), and a Prime 95B sCMOS camera (Photometrics), our setup employed illumination from a TTL triggered multifiber laser launch (Intelligent Imaging Innovations). For NRM’s in Figure 3, we used 405 nm and 561 nm lasers, while for NRVM’s in Figure 7, we employed a 561 nm laser. Imaging was performed using either a 63x, 1.4 NA Plan Apochromat or a 10x, 0.45 NA Plan Apochromat objectives (Zeiss). Temperature and humidity conditions were regulated using a Bold Line full enclosure incubator (Oko Labs). The microscope operations were controlled through Slidebook 6 Software (Intelligent Imaging Innovations). Images were processed using a Z-plane average picture of MyBP-HL and cMyBP-C channels. Then, cMyBP-C positive images were used to select sarcomere doublets and we determined regions of interest (ROI’s). Co-relation analysis was performed using Coloc2 Fiji plugin. Data are expressed as a correlation between cMyBP-C and MyBP-HL channels using a Pearson’s R index correlation coefficient.

### Peptide preparation

NRVMs were seeded at 1.5 million cells per well in a 6-well cell culture plate. NRVMs were transfected with 2.5 µg of full length MyBP-HL or MyBP-HL nonsense mutation plasmid as described in previous section. NRVMs were trypsinized using 0.25% Trypsin and collected in NRM maintenance media. Cells were spun down at 2000 x g for 4 mins, supernatant discarded, and cell pellets were washed with sterile cold DPBS, supernatant was discarded, and pellets were resuspended in 8 M Urea in 50 mM Ammonium Bicarbonate. Cell pellets were sonicated using Bioruptor Pico sonicator (Diagenode) and underwent 6 cycles of sonication, 15 sec on and 30 sec off, for 4 minutes and 30 seconds.

Protein samples were dissolved in 8 M urea in 50 mM ammonium bicarbonate (pH 7.8) by vortexing for 1 hour (Eppendorf ThermoMixer F1.5). 5 mM dithiothreitol (DTT; Sigma-Aldrich) was added to sample and incubated at 37 °C for 1 hour (Eppendorf ThermoMixer F1.5) to reduce disulfide bonds. To alkylate the samples, the samples were then incubated in 15 mM iodoacetamide (Sigma-Aldrich) for 30 minutes at room temperature. To quench the excess iodoacetamide, 25 mM DTT was added to samples and incubated at room temperature for 15 minutes. Protein samples were then diluted 1:4 with 50 mM ammonium bicarbonate (pH 7.8) to reduce the urea concentration. Samples were then digested and incubated with Mass Spectrometry Grade Proteases (Thermo Fisher) (1:50 wt/wt) overnight at 37 °C. Peptides were then purified and desalted using C18 spin columns (G-Biosciences), speed-vacuumed dry, and stored at -80 °C until further use. To determine peptide quantification, peptides were reconstituted in 0.1% formic acid and used in the Pierce Quantitative Colormetric Peptide Quantification Kit (Thermo Scientific).

### Mass Spectrometry

1.0 µg of resolubilized peptides were loaded onto a Vanquish Neo UHPLC system (Thermo Fisher) with a heated trap and elute workflow (c18 PrepMap, 5 mm, trap column, P/N 160454) in a forward-flush configuration connected to a 25 cm Easyspray analytical column (P/N ES802A rev2) 2 µM, 100 A, 75 µm x 25 with 100% buffer A (0.1% Formic acid in water) and the column oven operating at 40 °C. Peptides were eluted over a 150 minute gradient using 80% acetonitrile, and 0.1% formic acid (buffer B), going from 4% to 15% over 10 min, to 40% over the next 90 min, then to 55 % over the next 37 min, then to 99% over 6 min, and then kept at 99% for 7 min, after which all peptides were eluted. Spectra were acquired with an Orbitrap Eclipse Tribrid mass spectrometer with FAIMS Pro interface (ThermoFisher) running Tune 3.5 and Xcalibur 4.5. For all acquisition methods, spray voltage set to 2000 V, and ion transfer tube temperature set at 300 °C, FAIMS switched between compensation voltages (CVs) of -45 V, -55 V, and -65 V with cycle times of 1.5 seconds. MS1 spectra were acquired at 120,000 resolutions with a scan range from 375 to 1600 m/z, normalized AGC target of 300%, and maximum injection time of 50 ms, S-lens RF level set to 30, without source fragmentation and datatype positive and profile; Precursors were filtered using monoisotopic peak determination set to peptide; included charge states, 2 – 7 (reject unassigned); dynamic exclusion enabled, with n = 1 for 60 sec exclusion duration at ten ppm for high and low. DDMS2 scan used: isolation mode: quadrupole; isolation window: 1.6 m/z; activation type: HCD with 30% collision energy; IonTrap detector scan rate: turbo; AGC target: 10,000; maximum injection time: 35 ms; micro scans: 1, and data type: centroid.

### MS/MS data analysis

Raw data were analyzed using Proteome Discoverer 2.5 (ThermoFisher) using Sequest HT search engines. The data were searched against the *Mus musculus* Uniprot protein sequence database (UP000000589). The search parameters included precursor mass tolerance of 10 ppm and 0.06 Da for fragments, allowing two missed trypsin cleavages, oxidation (Met) and acetylation (protein N-term), and phosphorylation (S, T, Y) as variable modifications, and carbamidomethylation (Cys) as a static modification. Percolator PSM validation was used with the following parameters: strict false discover rate (FDR) of 0.01, relaxed FDR of 0.05, maximum ΔCn of 0.05, and validation based on q-value. Precursor ions quantifier settings were to use unique + razor for peptides; precursor abundance was based on intensity, normalization based on total peptide amount, protein abundance was calculated by the summed intensity of connected peptides, and protein ratios were calculated based on protein abundance and a background-based T-test was used to calculate statistical significance.

### Calcium transients

Neonatal ventricular myocytes were isolated, plated on GelTrex coated glass coverslips (22×22 mm; Corning) inside 60 mm culture plates and after 24 hours co-transfected with GFP alone and MyBP-HL nonsense mutations. Twenty-four hours after transfection, cells were rinsed with Tyrode’s buffer and incubated for 13 – 15 minutes in Tyrode’s buffer with 1 μM Fura-2 AM (ThermoFisher) at room temperature. After incubation, coverslips were refreshed with Tyrode’s free of Fura-2 AM for an additional 15 minutes at RT and then transferred to a recording chamber. NRVMs were visualized using a microscope (Nikon Instruments Inc) equipped with a MyoCam-S camera. GFP positive cells were first located, and all recordings were performed under electrical stimulation at 1 Hz for 5 ms and 35 – 40 V (IonOptix MyoPacer EP—Field Stimulator). The range of fluorescence emission was 340 – 380 nm and collected by a photomultiplier tube via the 40x objective during continuous excitation at 510 nm with a 75 W Xenon lamp. The IonOptix contractility system was used to quantify calcium transients and data was processed using IonWizard. Calcium transients were averaged and plotted using Prism8 software (GraphPad, San Diego, CA). We used at least 15 cells from five different NRM isolations. We analyzed and plotted the area under the curve of transients, FF, ms and Max_+df/dt_ Ca^+2^ (FF/ms).

### Contraction and sarcomere length measurements

To analyze if nonsense mutation were related with sarcomere contraction or sarcomere length anomalies in NRM’s, 2 x 10^6^ myocytes were plated on 22 x 22 glass coverslips (ThermoFisher, Cat. No: 22-050-218) that were coated with GelTrex matrix (ThermoFisher, Cat. No: A1413202). Cells were transfected with 2.5 mg of α-actinin-RFP (Addgene Cat. No: 58050). Pictures were obtained using a spinning-disk Zeiss 3i microscope at 25 frames per second for 10 seconds. Images were converted to 8 bits and transformed to AVI using Fiji software. Videos were sectioned to 150 frames to visualize the contraction and analyzed using python based Sarc-Graph automated segmentation, tracking and sarcomere analysis software (Zhao et al., 2021).

## Results

### Modeling *MYBPHL* nonsense mutations in NRVMs and NRAMs

We identified eight human premature stop mutations in *MYBPHL* in the GnomAD v2.1.1 database that we used in this work. We made single amino acid mutations in the mouse MyBP-HL sequence to introduce stop codons in the Q29, Trp54, W113, Y123, W158, W192, K250, and R255 codons (**Table 1**). We modeled these eight *MYBPHL* mutations in the mouse *Mybphl* cDNA sequence with an N’-terminal Myc-DDK tag (**Fig 1 A**). The anti-mouse MyBP-HL antibody does not recognize rat MyBP-HL, so we are able to transfect neonatal rat atrial cardiomyocytes (NRAMs) and differentiate our constructs from the endogenous rat MyBP-HL (**Fig. 1 B**). We tested these constructs by transfecting them into HEK293T cells and immunoblotting the total protein lysate (**Fig. 1 C**). The two smallest constructs did not appear on the immunoblots, so we re-probed the blot after removing the strong K250X and R255X samples (**Fig. 1 D**).

**Fig 1.**
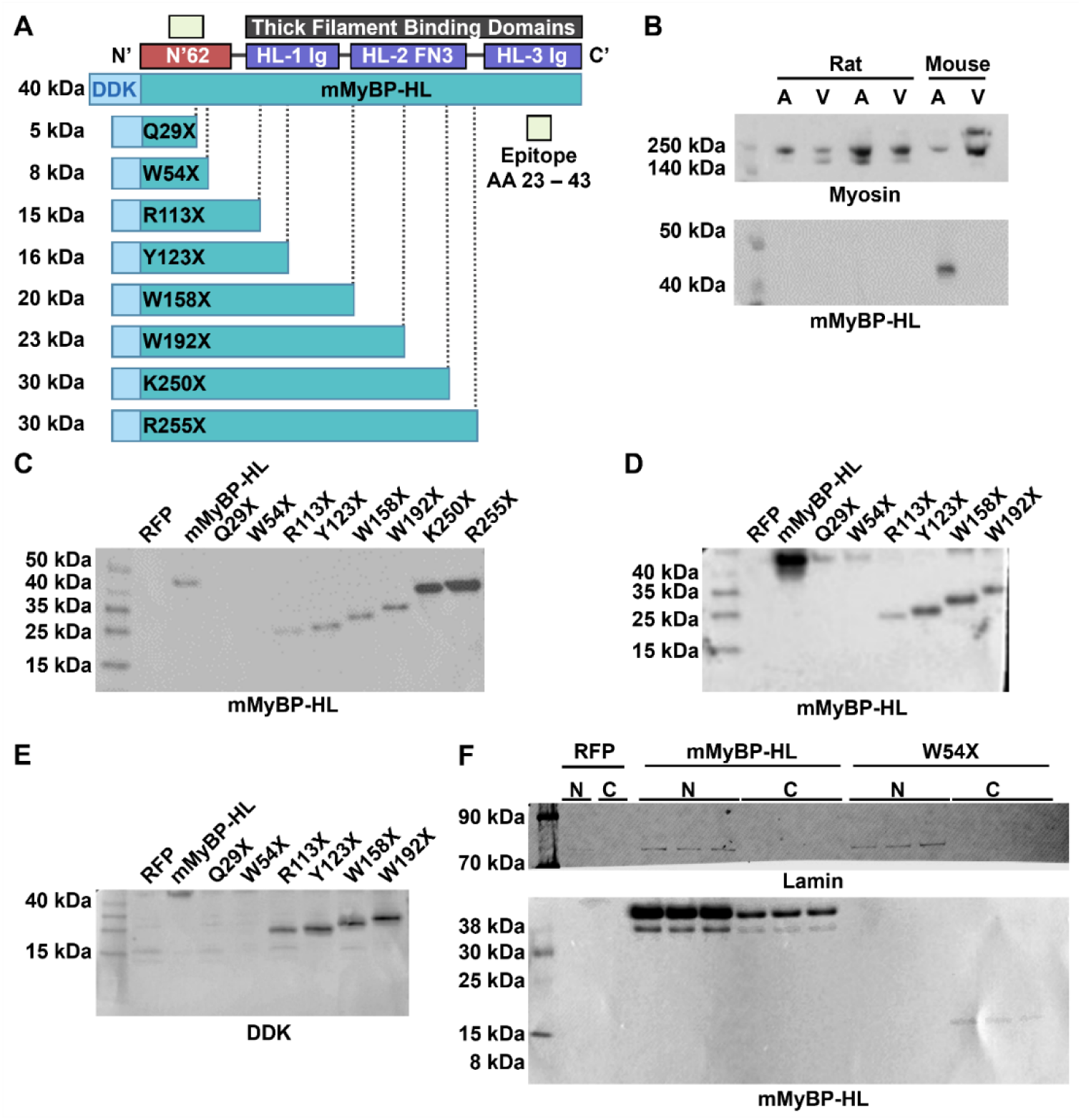
MyBP-HL nonsense mutations. **(A)** Schematic of the mouse MyBP-HL protein domains with the *Mybphl* cDNA containing an N’-terminal Myc-DDK tag below. Human missense mutations from gnomAD are shown below with the relative location of each truncating mutant. The epitope of the MyBP-HL specific antibody is marked in yellow. **(B)** Total protein from rat and mouse atria (A) and ventricle (V) samples immunoblotted for myosin heavy chain and mouse MyBP-HL. The MyBP-HL antibody does not detect rat MyBP-HL in the rat atria. **(C)** Immunoblot of HEK cell lysate after overexpression of *Mybphl* constructs to demonstrate the expected protein products. **(D)** Immunoblots were cut to remove the highly expressed K250 and R255 mutants to better visualize the smaller proteins. **(E)** We attempted to detect Q29X and W54X using the DDK epitope tag present in these constructs, but neither short truncation was detectable. **(F)** To identify Q29X and W54X, we resolved nuclear and cytoplasmic protein fractions on a low percent acrylamide gel and were able to detect W54X in the cytoplasm.

The epitope for the anti-mMyBP-HL antibody includes amino acids 21 – 40 (Barefield et al., 2022), and is therefore almost halved in the Q29X mutant, likely preventing detection. We immunoblotted using the N’-terminal DDK tag and again detected the full-length and longer truncated proteins, but not Q29X or W54X (**Fig. 1 E**). As the W54X mutant retains the complete antibody epitope, we also investigated whether W54X might be translated but sequestered in the nucleus. Fractionating protein into cytoplasmic and nuclear fractions did reveal W54X, although unexpectedly it was found in the cytoplasmic fraction, while Q29X was still not detected (**Fig. 1 F**). Finally, we performed mass spectrometry on NRVMs transfected with RFP, the nonsense mutations, and wild type MyBP-HL. The peptide closest to the N’-terminus found in the mass spectrometry data was 31-TSHQQEAGSPSLQLLPSIEEHPKIWLPR-58, downstream of the Q29X stop site. The inability to reliably detect Q29X led us to exclude it from further experiments.

NRAM’s and NRVM’s were then transfected with our constructs to identify their ability to incorporate into the sarcomere. To demonstrate that all the constructs overexpress at similar levels, we performed fluorometric experiments to evaluate the overall fluorescence of immunostained MyBP-HL and the MyBP-HL nonsense mutations. We normalized the overexpression values using the fluorometric values for endogenous actin marked with fluorescent phalloidin. MyBP-HL full length protein fluorescence was not significantly different than the MyBP-HL nonsense mutations, suggesting similar levels of overexpression for all the constructs (**Fig. 2**).

**Fig. 2.**
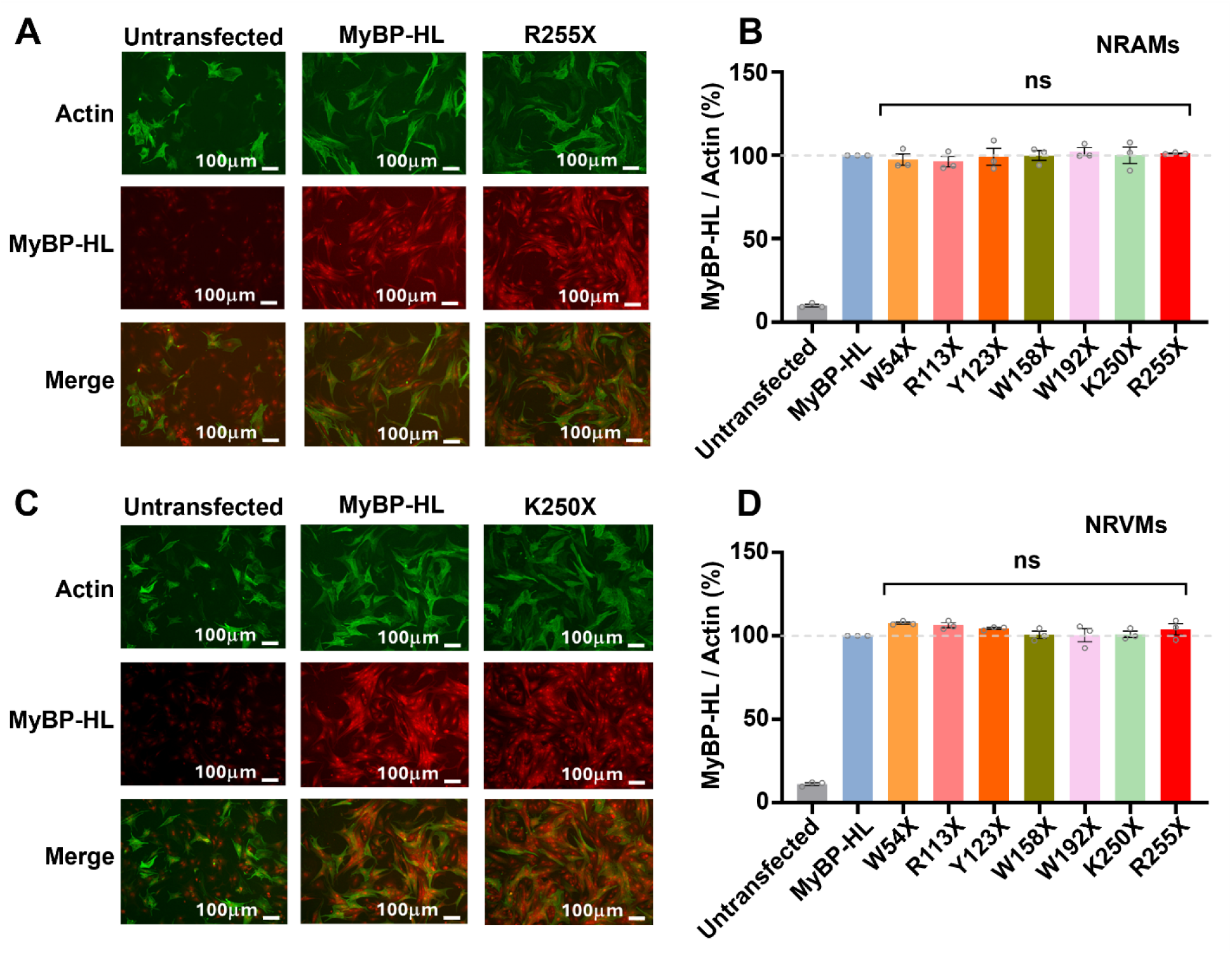
Overexpression levels are similar between full-length and nonsense mutations MyBP-HL in NRMs. **(A-B)** NRAMs and **(C-D)** NRVMs were transfected with MyBP-HL full length and MyBP-HL nonsense mutations. Cells were immunostained with MyBP-HL and actin specific antibodies. Alexa468 and Alexa568 were used as secondary antibodies, respectively. MyBP-HL fluorescence was measured and normalized using actin fluorescence. Results show similar levels of fluorescence for transfections with full-length MyBP-HL and MyBP-HL nonsense mutations. Results are expressed as the mean ± SEM of three independent experiments. Multiple comparison ANOVA and Dunnett test was performed compared to the positive control (MyBP-HL).

### MyBP-HL nonsense mutations disrupt MyBP-HL sarcomere localization

We next investigated the consequence of MyBP-HL nonsense mutations on the ability of MyBP-HL to incorporate into the sarcomere. We analyzed sarcomere incorporation through confocal microscopy. It’s known that MyBP-HL full length protein incorporates into the sarcomere where it co-localizes with cardiac myosin binding protein-C (cMyBP-C) in the C-zone doublets of the sarcomere (Barefield et al., 2023b) (**Fig. 3 B**). NRM’s were transfected with MyBP-HL full length and MyBP-HL nonsense mutations, then fixed and stained with MyBP-HL and cMyBP-C specific antibodies (**Fig 3 A, B**). We used NRM’s transfected with RFP as a negative control. We validated that full-length MyBP-HL properly incorporates into the sarcomere, forming C-zone doublets in a similar pattern to cMyBP-C that gave a 0.8 MyBP-HL/cMyBP-C co-localization index (**Fig 3 C, D**). Nevertheless, all MyBP-HL nonsense mutations showed significantly reduced correlation indices, indicating that MyBP-HL nonsense peptides are not completely incorporating into the sarcomere. The truncated proteins mostly localized diffusely, with some variable sarcomere enrichment and puncta throughout the cytosol. We observed a reduction in colocalization to ∼50%, meaning some of the overexpressed nonsense protein may bind to the sarcomeres, or the overexpression causes some incidental sarcomere colocalization. We also observed the same reduced sarcomere incorporation of MyBP-HL nonsense mutants in both NRAM’s and NRVM’s. Interestingly, the only truncating mutant that showed any pronounced sarcomere stripe localization was W54X. This mutant contains most of the N’-terminal 62 amino acid unstructured domain of MyBP-HL (**Fig. 1 A**), and this finding suggests that this domain may have some interactions within the sarcomere.

**Figure 3.**
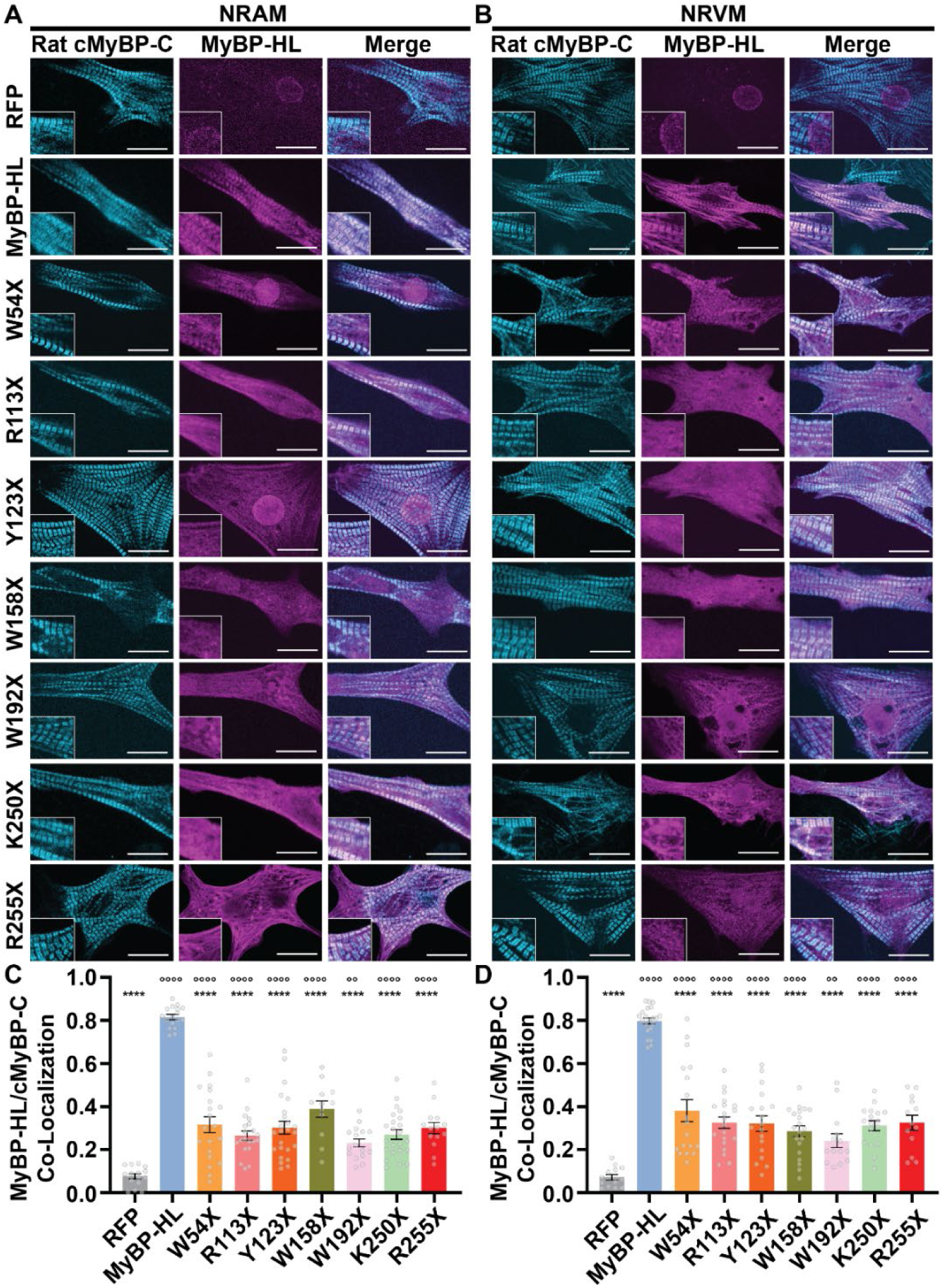
MyBP-HL nonsense mutations show improper sarcomere incorporation. **(A)** NRAM’s and **(C)** NRVM’s were transfected with MyBP-HL full length and MyBP-HL nonsense mutations. Cells were stained with specific antibodies for overexpressed mouse MyBP-HL and endogenous rat cMyBP-C. **(B and D)** Pearson’s co-localization test was performed to quantify MyBP-HL co-localization with endogenous rat cMyBP-C in the expected C-zone doublets. Results are expressed as the mean ± SEM. Multiple comparison ANOVA and Dunnett test was performed referred to cells transfected with RFP as a negative control.

### Overexpressed MyBP-HL is degraded by calpain and the ubiquitin-proteosome system

We next determined whether overexpressed MyBP-HL nonsense proteins are degraded differently than the full-length control. The consensus on nonsense mutations in the highly related cMyBP-C is that truncated mutant proteins are degraded rapidly and do not generate poison peptides (Barefield et al., 2022). Therefore, we aimed to determine if different MyBP-HL nonsense peptides are degraded differently. We performed computational analysis to identify possible degradation regions in MyBP-HL for calpain and the ubiquitin-proteasome system, both of which frequently degrade sarcomere proteins. We identified two calpain substrate regions in MyBP-HL at residues E18 and E166 and three ubiquitin-proteasome system processing regions at residues W88, W167, and Y200.

NRM’s were transfected with MyBP-HL full length and then incubated with 1 µM calpastatin and/or 100 nM epoxomicin for 24 hrs. We performed western blot analysis measuring MyBP-HL protein amount (**Fig. 4 A**). We used NRM’s transfected with RFP as a negative control. We showed that calpastatin and/or epoxomicin treatment increased abundance of MyBP-HL protein, suggesting that calpain and the ubiquitin-proteasome system can regulate degradation of overexpressed MyBP-HL. In cells treated with only calpain inhibitor (CI), we observed an increase of ∼90% of MyBP-HL. In cells treated with only proteasome inhibitor (PI) we observed an increase of ∼350% of MyBP-HL abundance. In cells treated with CI and PI together, we observed an increase of ∼380% of MyBP-HL abundance. These results show that calpain and the proteasome both regulate MyBP-HL degradation in a synergistic manner. Degradation results were similar in NRAM’s and NRVM’s transfected with MyBP-HL full length (**Fig. 4 A, B**), suggesting the competition with endogenous MyBP-HL and cMyBP-C for myosin binding sites is a negligible factor in an overexpression system. Importantly, we did not see a predominant proteolytic product of MyBP-HL when calpain was the sole protease, which we would expect as calpain usually makes specific degradation products (Barefield et al., 2019). This could be due to a predicted calpain site at E18 disrupting the antibody epitope site.

**Fig 4.**
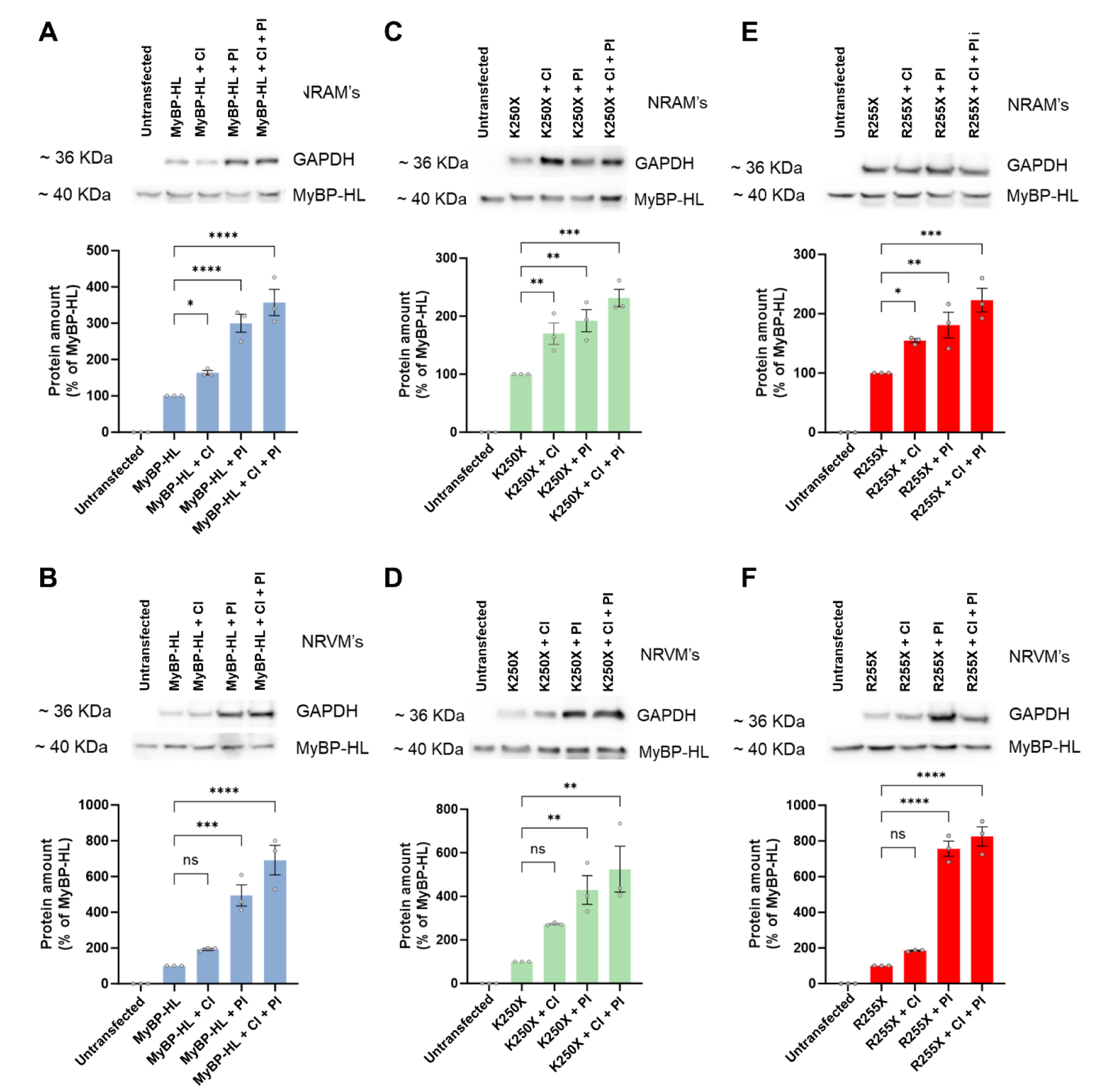
Calpain protease and the proteasome degrade full length and MyBP-HL nonsense mutations. WB analysis was performed to analyze the contribution of calpain and proteasome proteolysis on MyBP-HL full length and MyBP-HL nonsense mutations in NRAM’s and NRVM’s transfected with **(A)** MyBP-HL full length, **(B)** K250X, and **(C)** R255X. 24 hours after transfection cells were incubated with 1mM of calpastatin (CI) and 100 nM epoxomicin (PI) during 24 h. Results are expressed as the mean ± SEM of three independent experiments. Multiple comparison ANOVA and Holm-Sidaks test was performed referred to positive control (MyBP-HL).

We performed the same experiment using cells transfected with MyBP-HL nonsense mutations K250X and R255X (**Fig. 4 B, C**). The R255X mutation has a relatively high allele frequency, so we were interested in determining if this mutant had different degradation properties as a similarly sized nonsense mutation. We observed that degradation of K250X and R255X is facilitated by calpain and the proteasome with similar results as the full length MyBP-HL.

### MyBP-HL nonsense mutations promote aberrant calcium transients

We performed calcium transient experiments (**Fig. 5 A**) to assess whether MyBP-HL nonsense mutations alter cardiomyocyte function. To identify NRMs transfected with the MyBP-HL constructs, we co-transfected NRMs with the MyBP-HL constructs and a GFP reporter plasmid. Compared to full-length MyBP-HL, cells transfected with MyBP-HL nonsense mutations did not show significant differences in peak calcium (**Fig. 5B**), total amount of calcium released in each transient, which was calculated analyzing the area under the curve of the calcium transients (**Fig. 5C**), or time to peak calcium (**Fig. 5D**). These parameters suggest that overexpression of full-length MyBP-HL does not alter calcium handling, nor does overexpression of the nonsense mutants.

**Figure 5.**
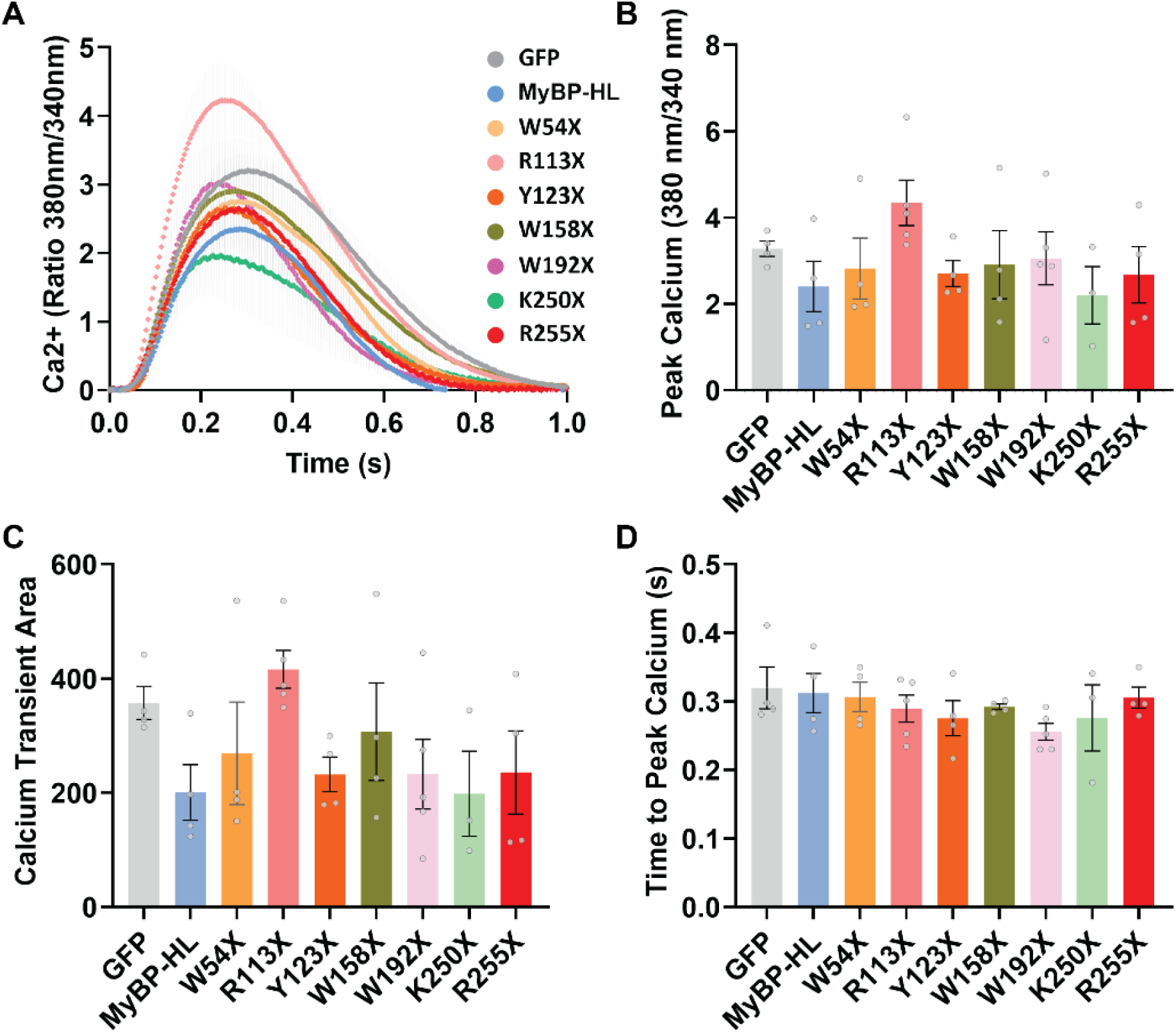
MyBP-HL nonsense mutations do not alter calcium transients. NRVM’s were transfected with MyBP-HL nonsense mutations and 24 hours later cells were incubated with 2 mM FURA2-AM. Transients were analyzed by fluorescence microscopy. **(A)** Calcium transient profiles of MyBP-HL full length and MyBP-HL nonsense mutations. **(B)** Peak calcium transient amplitude. **(C)** Total area under the curve of calcium transients showed in A. Results are expressed as the mean ± SEM of four independent experiments. **(D)** Time to peak calcium release at each cycle length shown in A. N=4 – 5 cell isolations, averages of 3 – 7 cells per isolation. One-way ANOVA was performed and compared against the MyBP-HL full-length.

### MyBP-HL nonsense mutations lead to altered contractility

We analyzed contractility of NRVM’s to examine any potential contractility alterations induced by MyBP-HL nonsense peptides. We co-transfected NRM’s with our MyBP-HL nonsense mutation plasmids and α-actinin-RFP plasmids. The α-actinin-RFP labeled the sarcomere Z-disks (**Fig. 6 A**). Using the α-actinin-RFP signal, we performed an automated quantitative analysis of NRM contraction in paced NRVMs using the python-based software Sarc-Graph (Zhao et al., 2021) (**Fig. 6 B**). Overexpression of full length MyBP-HL did not alter contractility compared with NRVM’s transfected with α-actinin-RFP alone. We did not observe changes in the number of beats per second between groups, showing that the cells were paced successfully at 1 Hz (**Fig. 6 C**). NRVM’s transfected with MyBP-HL nonsense mutations showed a significant reduction in contractile amplitude compared to cells transfected with full length MyBP-HL (**Fig. 6 D**). Interestingly, R255X did not show reduced contractile amplitude, with values like full length MyBP-HL, but different than the other nonsense mutations. With the aim to analyze changes in the sarcomere structure, we also used Sarc-Graph (Zhao et al., 2021) to analyze sarcomere length (**Fig. 6 E**). We observed a 0.2 µm statistically significant increase in sarcomere length for all MyBP-HL nonsense mutations (**Fig. 6 F**). Interestingly, we did not observe an increase in sarcomere length of cells transfected with MyBP-HL full length compared to the RFP control.

**Figure 6.**
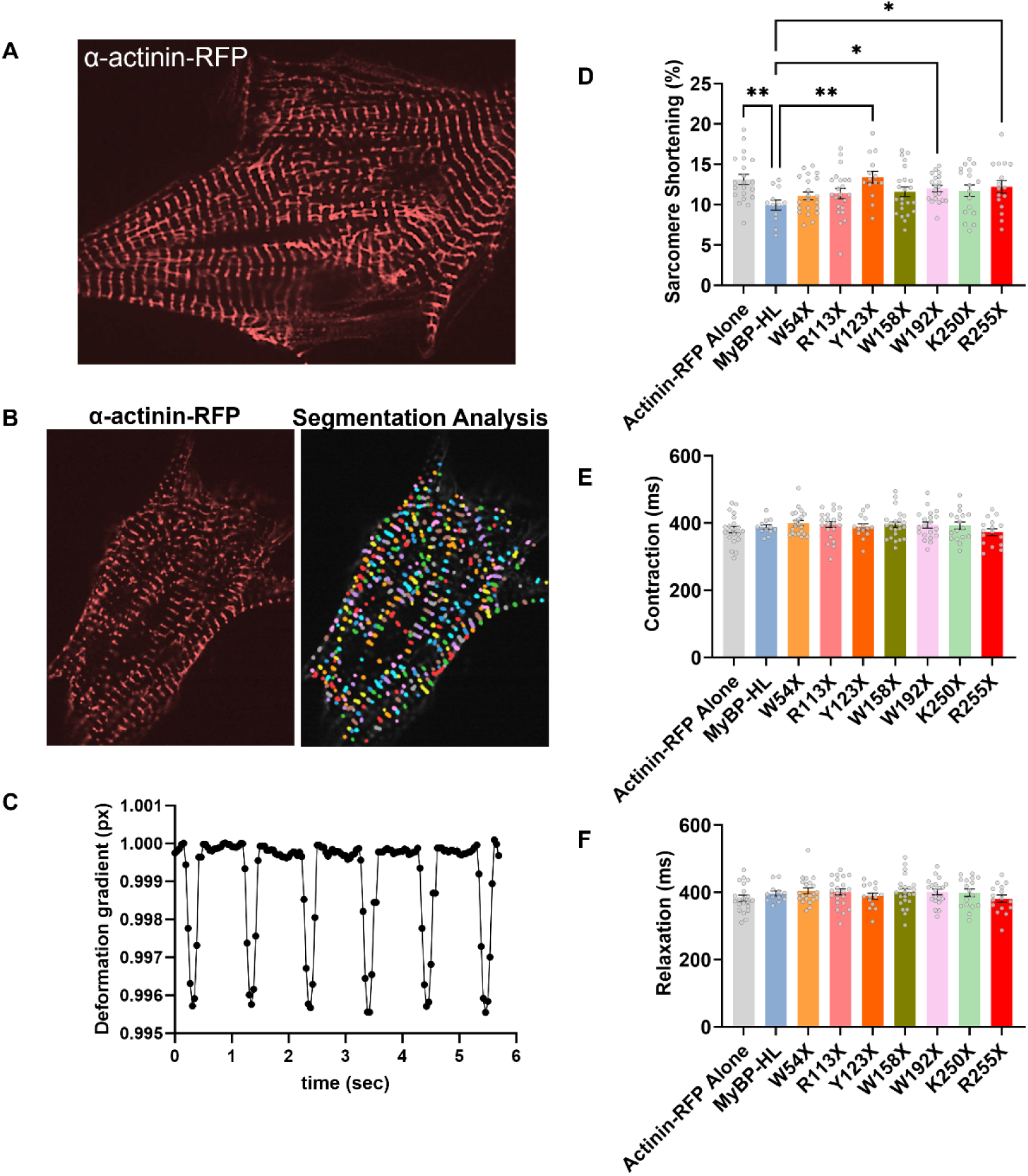
MyBP-HL nonsense mutations lead to a reduction in contraction amplitude and increase the sarcomere length. NRVM’s were co-transfected with α-actinin-RFP and MyBP-HL plasmids. **(A)** Videos were obtained by spinning disk confocal microscope. **(B)** Sarcomeres are automatically detected and segmented to allow tracking of sarcomere shortening during a contraction. **(C)** Deformation gradients were calculated using SARC-GRAPH software. **(D)** The percent shortening, **(E)** time to peak contraction, and **(F)** time from peak contraction to relaxation are displayed. N=3 cardiomyocyte isolations were performed with N=3 isolations per group, with 6 – 10 cells analyzed per sample. Results are expressed as the mean ± SEM. Multiple comparison ANOVA and Dunnett test was performed using MyBP-HL full-length as the repeated measures control.

## Discussion

The recent identification of the atrial-specific myosin binding protein MyBP-HL has revealed yet another difference between atrial and ventricular sarcomere regulation (Burnham et al., 2024). We have previously shown that MyBP-HL and cMyBP-C have an inverse relationship, where overexpression or knock-down of either protein causes an inverse change in the levels of the other protein (Barefield et al., 2023b). Because the levels of these two regulatory proteins is dictated by their ability to compete for sarcomere incorporation, nonsense mutations that remove the C’-terminal binding domains of either protein would cause an increase in the opposing myosin binding protein.

### Myosin binding protein nonsense mutations

The initial identification of MyBP-HL as a DCM-linked gene was from a family that carried the *MYBPHL* R255X mutation that is missing the HL-3 C’-terminal domain (Barefield et al., 2017). However, that R255X mutation is potentially present in ∼120 million people in the population. This makes it unlikely to be a fully penetrant autosomal dominant DCM-causing mutation, although our understanding of how atrial dysfunction contributes to ventricular myopathy is emerging (Burnham et al., 2024). To better understand the R255X and other *MYBPHL* nonsense mutations, we now analyzed the implications and physiological consequences of the expression of MyBP-HL nonsense mutations to determine if the R255X mutation behaved differently than the others. The 12 *MYBPHL* nonsense mutations were selected from GnomAD v.2.1.1, which were all the premature stop mutations listed. Since then, GnomAD v4.0.0 now reports 21 premature stop variants in *MYBPHL*. As we saw highly consistent changes between most of our tested mutations, and those premature stops spanned the length of the gene, we expected any of the new untested mutations would behave similarly.

We found that the sarcomere incorporation of MyBP-HL nonsense mutations was overall lower, but highly variable. Overexpression of full length MyBP-HL correlated highly with cMyBP-C, indicating a proper sarcomere incorporation as these two proteins are known to both occupy the C-zones of the sarcomere (Barefield et al., 2023b). Based on the work on cMyBP-C truncating mutations, we expect that loss of even half the HL-3 domain would cause failure in sarcomere incorporation (Welikson and Fischman, 2002). However, we observed that the W54X mutation, lacking the HL1-Ig, HL2-FN3, and HL3-Ig domains, maintains some ability to localize to the sarcomere. These findings are important, as MyBP-HL has a unique N-terminal 62-amino acid domain that has no structural homology to other known protein domains. We hypothesize that this N’-terminal domain may be able to interact with components of the sarcomere, although a large systematic study will be required to test this hypothesis.

Nonsense mutations in *MYBPC3* are known to be rapidly degraded, likely by nonsense mediated decay of the mRNA. This illustrates a limitation in our study, as we are forcing the overexpression of these *MYBPHL* nonsense mutations. However, these comparisons are still informative of the necessity of specific protein domains for sarcomere incorporation of MyBP-HL.

### Proteolysis of sarcomere proteins

Sarcomere proteins are known to undergo degradation through calpain proteolysis and the ubiquitin proteasome system, and these processes are involved with or altered by disease states (Schlossarek et al., 2014; Barefield et al., 2019; Martin and Kirk, 2020).

We expected to observe the participation of the proteasome and calpain in degradation of the overexpressed proteins, but we were unsure whether MyBP-HL nonsense mutations would be degraded by these processes differently than full length MyBP-HL. We observed that both calpain and the proteasome degrade MyBP-HL full length and MyBP-HL nonsense mutations constructs.

### Functional changes

We observed that overexpression of these MyBP-HL constructs did not significantly alter calcium handling. This suggests that overexpression of the *MYBPHL* nonsense mutants impairs contractility, although we did not detect any overt differences in cardiomyocyte or sarcomere morphology between wild type and nonsense transfected cells. The analysis of the physiological implications of the overexpression of the MyBP-HL nonsense mutations resulted in a reduction in the amplitude of the contraction patterns analyzed and interestingly an increase of sarcomere length.

### Limitations

While our study provides valuable data, the interpretation of the results should consider the inherent limitations of overexpression studies. *MYBPC3* nonsense mutations are known to undergo rapid degradation and do not naturally build up protein aggregates (Barefield, 2023). Our study primarily utilizes MyBP-HL overexpression experiments, which can result in protein aggregation and non-specific effects, potentially influencing the outcomes. To mitigate this issue, we employed full-length MyBP-HL as a control. Although this approach reduces the likelihood of artifacts, some findings may still be affected by the elevated protein expression levels. Further validation using cell or animal models with *Mybphl* constructs knocked-in to the endogenous locus is recommended to corroborate these results. These comparisons are still valuable for elucidating the essential protein functions of MyBP-HL within the sarcomere.

### Conclusion

We have confirmed that MyBP-HL nonsense mutations cause improper sarcomere incorporation. We demonstrated that the degradation of overexpressed full length MyBP-HL and MyBP-HL nonsense mutations occurs through the proteasome as we expected, but also through the activation of calpain. We found that the MyBP-HL nonsense mutations lead to an increase in total calcium release during calcium transients. These phenomena seem to be related to changes in the contraction of NRM’s. The alterations in MyBP-HL nonsense peptides to sarcomere incorporation, degradation, and calcium transients, leads to a deficiency in contractility, although extensive work will need to be done to identify the mechanism of these dysregulations.

## Abbreviations

(DCM): Dilated Cardiomyopathy
(AF): Atrial fibrillation
(MyBP-HL): Myosin-binding protein-H like
(MyBP-HL-FL): Myosin-binding protein-H like full length
(MyBP-HL-ns): Myosin-binding protein-H like nonsense mutations
(cMyBP-C): Cardiac myosin binding protein-C

## Author contributions

A.A.A. and D.Y.B. conceived and designed research; A.A.A., G.E.F., H.V.B., K.N.A., A.P., and L.W. performed experiments; A.A.A., H.V.B., K.N.A., and D.Y.B. analyzed data; A.A.A., H.V.B., K.N.A., and D.Y.B. interpreted results of experiments; A.A.A. and D.Y.B. prepared figures; A.A.A. and D.Y.B. drafted the manuscript; A.A.A., G.E.F., H.V.B., K.N.A., and D.Y.B. edited and revised the manuscript; all authors approved the final version of the manuscript.

